# Perdurant TTC21B protein in the early mouse embryo is required for proper forebrain neural progenitor proliferation

**DOI:** 10.1101/2025.01.14.632919

**Authors:** Rebekah Niewoehner, David Paulding, Jesus Leal, Rolf W. Stottmann

## Abstract

Primary cilia play a pivotal role in cellular signaling and development and disruptions in ciliary form and/or function leads to human ciliopathies. Here, we examine the role of *Ttc21b*, a key component of the intraflagellar transport-A complex, in mouse forebrain development using a *Ttc21b*^*alien*^ null allele. Our findings reveal significant microcephaly in homozygous mutants is caused by disrupted neural progenitor proliferation and differentiation. Histological and immunohistochemical analyses show an enlarged ventricular zone and reduced cortical plate thickness, accompanied by altered mitotic spindle angles, suggesting defects in symmetric versus asymmetric cell divisions. Despite low *Ttc21b* expression in the forebrain epithelium, early embryonic expression patterns imply that perdurant TTC21B protein may underlie these phenotypes. Progenitor proliferation kinetics were disrupted, with fewer cells re-entering the cell cycle, correlating with reduced TBR2-positive intermediate progenitors and altered neurogenesis dynamics. Neuronal processes in the cortical plate were significantly shortened, suggesting cytoskeletal defects specific to terminal differentiation stages. Our findings support a model where early *Ttc21b* expression in precursors destined for the forebrain is critical for sustaining later neural progenitor proliferation and differentiation. These results advance our understanding of primary cilia in cortical development and provide a framework for exploring cytoskeletal contributions to ciliopathies.

## INTRODUCTION

The primary cilium is a microtubule-based extension of the cell found on virtually every cell type. These organelles have recovered in the past three decades from being called vestigial structures (Alberts 1994) and are now known to be crucial for proper signal transduction in several contexts. Primary cilia are supported by the axonemal microtubules anchored to the basal body/centriole.

Several different signal transduction effector molecules require trafficking through the cilium mediated by evolutionarily conserved intraflagellar transport (IFT) proteins, most notably probably the GLI proteins required for hedgehog signaling. Kinesin motors move IFT-B complexes and associated cargo along the microtubules of the axoneme towards the distal tip in an anterograde manner, and dynein motors move IFT-A complexes and cargo in a retrograde fashion back towards the cell body. IFT-A transport is also essential for organizing material at the base of cilia, especially GPCRs (Fig 1) (Baker and Beales 2009; Goetz and Anderson 2010; Reiter and Leroux 2017; Wheway et al. 2018). In mammalian systems, the primary cilium is required for proper regulation of multiple signaling pathways inside the cell, including the *Hedgehog*, polycystin, *Notch, BMP, PDGF*, GPCR, Hippo, mTor, canonical *Wnt*, and planar cell polarity pathways(Goetz and Anderson 2010; Wheway et al. 2018). Primary cilia have been previously shown to be crucial for forebrain development in multiple contexts (Zaidi et al. 2022).

**Figure 1.**
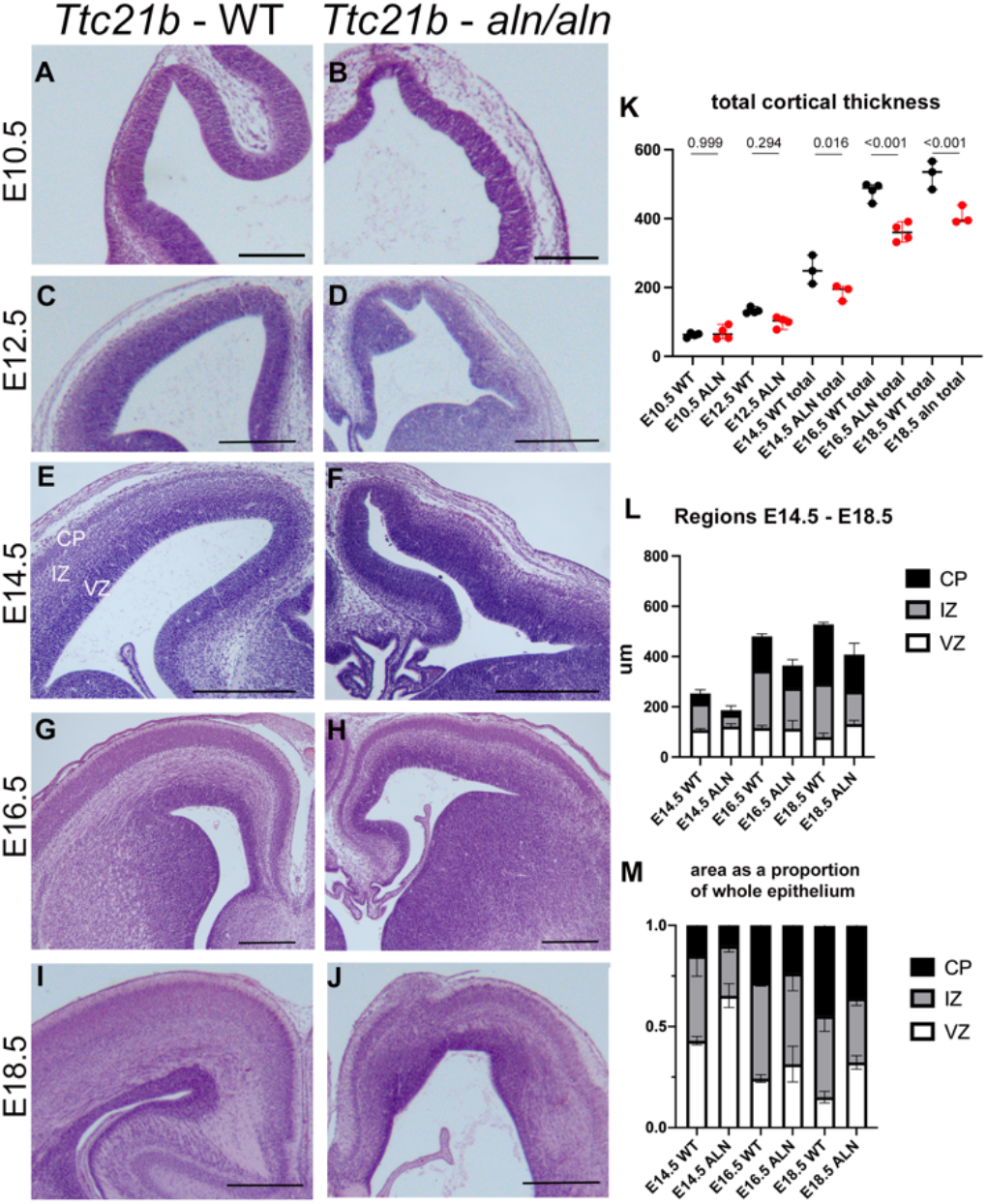
Histological analysis of *Ttc21b*^*alien*^ cortical development. Sections of wild-type (A,C,E,G,I) and *Ttc21b*^*aln/aln*^ (B,D,F,H,J) brains at E10.5 (A,B), E12.5 (C,D), E14.5 (E,F), E16.5 (G,H), and E18.5 (I,J). Total cortical thickness is quantified (K, t-test p values shown) and shows robust reduction in mutants by E14.5. Width of the ventricular zone (VZ), intermediate zone (IZ), and cortical plate (CP) are compared between wild-type (WT) and *Ttc21b*^*aln/aln*^ (ALN) brains (L) and shown as proportions of total width (M). Scale bars in A,C,E = 500 µm, G,I = 1mm.

*Tetratricopeptide repeat domain 21B* (*Ttc21b/Ift139*) is a component of the intraflagellar transport-A complex and a null allele of *Ttc21b* recovered from an ENU mutagenesis forward genetic screen has multiple embryonic phenotypes consistent with ciliopathies (Tran et al. 2008).

We have previously shown that a null allele of *Ttc21b* leads to a smaller forebrain and an anterior-posterior patterning phenotype in the early developing anterior nervous system (Stottmann et al. 2009). We attempted to use conditional deletion of *Ttc21b* in the developing forebrain to study the molecular basis of this microcephaly in the absence of any confounding effects of the patterning deficit. Much to our surprise, deletion of *Ttc21b* using the established *Foxg1-Cre* and *Emx1-Cre* tools did not recapitulate the microcephaly phenotype seen in the germline null animals (Snedeker et al.2017). The control of neural progenitor proliferation is thought to be autonomous to the forebrain tissue, making these phenotypes particularly intriguing. We have continued to study the processes leading to microcephaly in the *Ttc21b* mutants to elaborate on the role of primary cilia in neural progenitors and forebrain development.

## MATERIALS AND METHODS

### Mouse husbandry

All animals were maintained through a protocol approved by Nationwide Children’s Hospital Medical Center IACUC committee (IACUC2021-AR1200067). Mice were housed with a 12-h light cycle with food and water *ad libitum*. Mouse euthanasia was performed in a carbon dioxide chamber followed by secondary cervical dislocation. Genotyping was performed via PCR and gel electrophoresis on a 2% agarose gel or custom Taqman assays. All mouse alleles are previously published. Whole-brain and skeletal images were taken on a Zeiss Discovery V12 microscope (Zeiss, St. Louis, MO).

### Histology

Brains were dissected, fixed in formalin for 48h, washed in 70% ethanol, then dehydrated and paraffin-embedded by the morphology core. Blocks were sectioned on a microtome at 10µm (Sakura, Hayward, CA). Sections were placed on glass slides (Cardinal Health, Dublin, OH), baked >1 hour, and stained with hematoxylin and eosin using standard methods. Body and brain weights were obtained on a standard chemical scale. For cortical and cerebellar measurements, area in µm^2^ or length in µm was measured on ZEN 3.7 software. A minimum of 3 animals from at least 2 distinct litters were measured for each genotype. Embryos were stained for lacZ using standard protocols (Behringer 2014).

### Anatomical measurements

Matched sections from control and mutant embryos were chosen to be comparable in anterior-posterior position across the ages E10.5, E12.5, E14.5, E16.5, and E18.5. The cortical thickness was measured at the central-most point of the right dorsal pallium of the cortex. Measurements were taken using ImageJ for total cortical thickness, averaged, and compared for controls and mutants of all stages. For the stages E14.5, E16.5, and E18.5, the cortex was divided into Ventricular Zone, Intermediate Zone, and Cortical Plate. This separation was determined by examining morphology by eye, and was kept consistent across all samples.

### Immunohistochemistry

After dissection, brains were fixed for 1-2 days in PFA at 4°C. PFA was replaced with 30% Sucrose for 2 days before brains were embedded in Optimal Cutting Temperature solution (Sakura) and stored at -80°C. 10µm sections were obtained for mouse brain and human organoid samples on a Leica CM 1860 cryostat, placed on glass slides, and stored at - 20°C. Slides selected for IHC were pre-warmed at 42°C for 10-15 minutes.

Antibody retrieval was performed as previously described (Bittermann et al., 2019). The following antibodies were used: pHH3 (Sigma Aldrich AB_477043, 1:500), TBR2 (Abcam ab23345, 1:200), CC3 (Cell Signaling #9661, 1:300), gamma tubulin (Sigma T6557, 1:1000),

Alexa Fluor 488 Goat anti-Rabbit (InvitrogenA11008, 1:500), Alexa Fluor 488 Goat anti-Mouse (Invitrogen A11001, 1:500), Alexa Fluor 610 Goat anti-Rabbit (Invitrogen A20980, 1:500), Alexa Fluor 555 Goat anti-Mouse (InvitrogenA21442, 1:500). Slides were mounted in Prolong Gold Antifade medium (InvitrogenP36935). The MORF3 alleles allows staining with a V5 antibody (Invitrogen R960-25, 1:1000).

### MORF3 visualization

Brains were sectioned 40 µm on a Leica CM 1860 cryostat, placed on glass slides, and stored at -20°C. Slides selected for immunohistochemistry were pre-warmed at 42°C for 10-15 minutes. Antibody retrieval was performed as previously described (Bittermann et al, 2019). Confocal imaging for the MORF3 allele was done with a Nikon AX R Confocal microscope and analyzed using NIS Elements. For the quantification of neuronal morphology in the cortex, neurons with three or more processes were considered multipolar and neurons with two processes were considered bipolar.

*EdU/Ki67 staining and quantification* EdU (Invitrogen) was injected at a concentration of 20mg/kg interperitoneally into pregnant females. Embryos were harvested 24 or 72 hours later and stained as above. Ki67 primary antibody (1:200 Abcam ab15580) was added overnight at 4°C. A goat anti-rabbit secondary antibody (1:500 Invitrogen Alexa-Fluor 594) was added for 1 hour at room temperature. Slides were washed twice with 1X PBS and once with 3% BSA in PBS. EdU was labeled using the Click-iT EdU Alexa Fluor 488 Imaging Kit (Invitrogen). Quantification was done using Imaris software package. A surface was manually created that contained the cortex from the VZ to the CP and the enclosed area was quantified. EdU positive cells and Ki67 positive cells were then identified using the Imaris “Spots” function. Cells both EdU and Ki67 positive were quantified using the “Colocalize Spots” function. Each positive cell count was normalized to the area of the section.

### Statistical analysis

Data plots and subsequent analyses were performed with Prism 9 (GraphPad, San Diego, CA). A student’s t-test was performed for experiments with two groups. An ANOVA with Tukey’s multiple comparison tests was performed for experiments with more than two comparisons. ANOVA p-values are usually stated in the text or figure legend and specific relevant p-values for the multiple comparisons are shown in the relevant figure. We report the statistical test values directly rather than assigning a significance symbol to provide all the data for the reader. Data shown are the median +/-95% confidence interval.

## RESULTS

### Ttc21b-aln brains are significantly smaller with disruptions to the proliferative ventricular zone

Previous work has shown that conditional deletion of *Ttc21b* from the forebrain did not result in smaller brain size as is true for *Ttc21b*^*alien/alien*^ homozygous mutants (Snedeker et al. 2017). In order to understand the mechanisms leading to such a reduction, we returned to analyze the null allele. We have previously shown that *Ttc21b*^*alien/alien*^ homozygous mutants on an FVB genetic background have an even more severe phenotype than when maintained on a largely B6 background (Snedeker et al. 2019). However, other previous work from our group has shown that congenic FVB mice lead to high rates of exencephaly and/or embryonic death preventing an efficient study of forebrain development. We therefore performed this study reported here only on embryos derived from intercrosses of *Ttc21b*^*alien/wt*^ heterozygous carriers maintained on an FVB with those maintained on a B6 background: F1 hybrids. We previously showed that the forebrains of *Ttc21b* ^*alien/alien*^ mice are appreciably smaller as early as E12.5 (Stottmann et al. 2009), Fig. 1). Here we examine the cortical epithelium from embryonic day (E) 10.5 through E18.5. Histological features of *Ttc21b*^*alien/alien*^ mutants include an irregular edge to the epithelium in mutants at E10.5 (Fig. 1B). The size of the proliferative ventricular zone appears larger as soon as it is a distinctive layer within the cortex while the cortical plate housing differentiated neurons appears smaller (Fig. 1F, H, J). We note that the differences in mutants described here are somewhat variable, even within a mutant brain, and throughout the totality of the mutants studied as a whole. We quantified the width of the cortex from E10.5 through E18.5, as well as the ventricular zone, intermediate zone and cortical plate from E14.5-E18.5. While obviously dysmorphic and irregular at E10.5, the total cortical thickness is significantly reduced by E14.5 and only further deviates over time (Fig. 1K). While the cortex is thinner, the relative proportion of the area taken up by the ventricular zone is greater at all stages examined as the cortical plate is always smaller (Fig. 1L,M).

### Ttc21b expression is not high in forebrain

The surprising results from the genetic deletions of *Ttc21b* from the forebrain (Snedeker et al. 2017) motivated us to take a much closer look at the expression of *Ttc21b* throughout forebrain development. We used the lacZ expression from the *Ttc21b* ^*tm1a*^ gene trap allele and see high expression throughout the embryo at E7.5 (Fig. 2A) and E8.5 (Fig. 2B). By E9.5 (Fig. 2C) and E10.5 (Fig. 2D), whole mount analysis clearly shows that *Ttc21b* expression becomes much more restricted, but the dorsal forebrain is an area that appears to retain lacZ signal. Histological section analysis shows the expression is most prominent in the layer immediately adjacent to the neural epithelium (Fig. 2E,F, arrow in F), which is most likely the neural-crest derived meningeal layer. Close examination does reveal scattered lacZ-positive cells within the forebrain neural epithelium, but we see no clear pattern to these cells (Fig. 2G arrows). Whole mount analysis at E12.5 indicates even lower *Ttc21b* expression and sections again show that there are very few positive cells scattered throughout the epithelium. We note they are again much more obvious in the layer of cells immediately adjacent to the epithelium (Fig. 2I-K). Sections at E14.5 show this pattern continues but we also note a new population of cells in the intermediate zone (Fig. 2M, arrow). Given the previously demonstrated role of *Arl13b* in tangential migration (Higginbotham et al. 2012) and the position of these cells, we hypothesize these are interneurons migrating from the ventral brain structures towards their final destination in the cortical plate. We note interesting expression in increasingly later stages of brain development but these are not immediately relevant to the rest of the work presented here.

**Figure 2.**
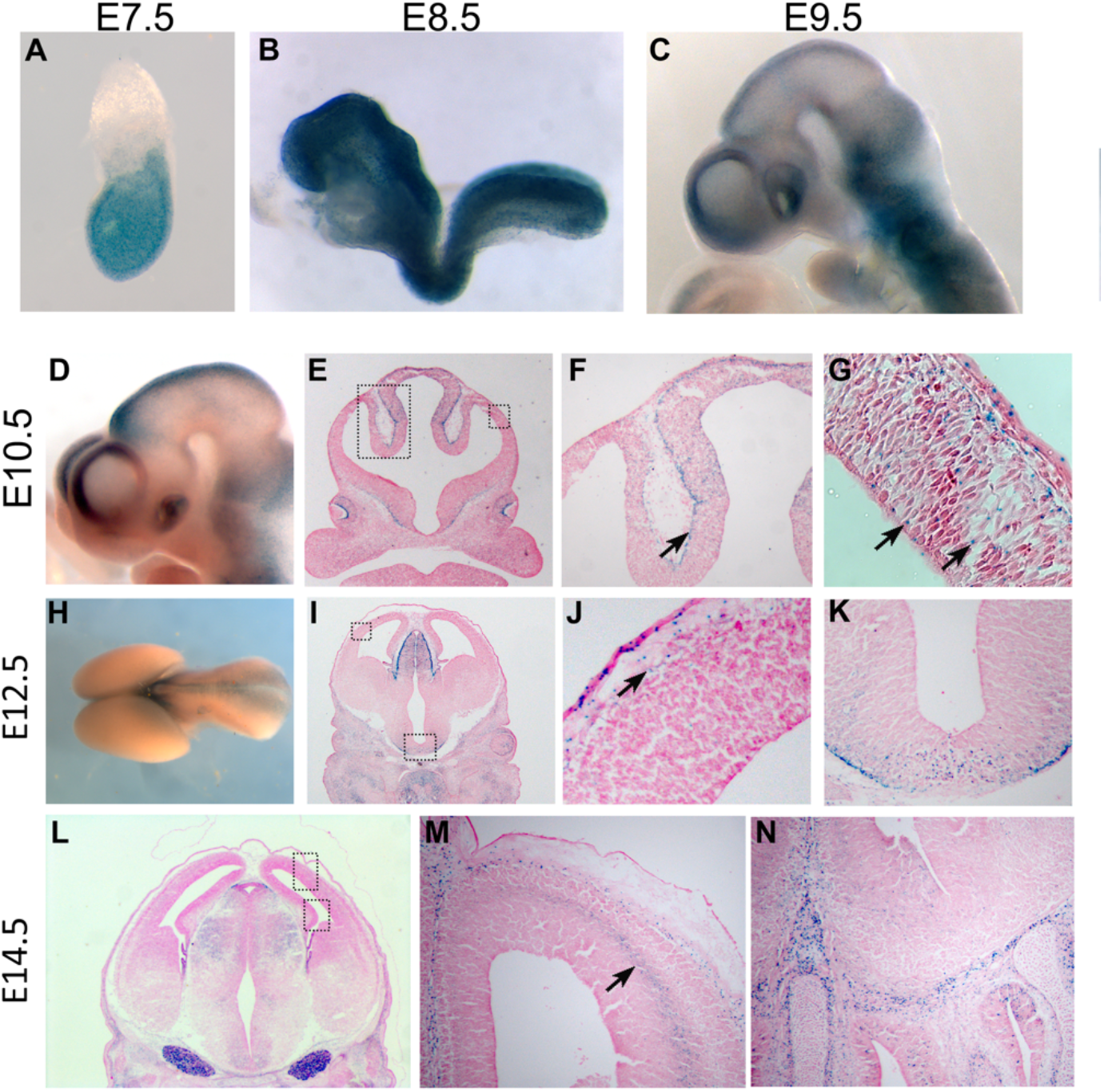
*Ttc21b* lacz expression. *Ttc21b*^*tm1a*^ embryos were stained with X-gal to highlight *Ttc21b* expression in whole mount embryos (A-D, H) or after sectioning (E-G, I-N) at E7.5 (A), E8.5 (B), E9,5 (C), E10.5 (D-G), E12.5 (H-K), and E14.5 (L-N). Boxes in E show areas highlighted in F and G, boxes in I and L show areas magnified in J and M, respectively.

These findings determining the expression patterns of *Ttc21b* may explain our previous results in which *Foxg1-Cre* and *Emx1-Cre* did not recapitulate the *Ttc21b*^*aln*^germ line phenotype (Snedeker et al. 2017). These Cre transgenes drive Cre recombination in the neural epithelium which is clearly not a region of high *Ttc21b* expression (Hebert and McConnell 2000; Gorski et al. 2002). Rather, the highest levels of *Ttc21b* gene expression are at earlier stages in tissues which will in turn give rise to the cortical epithelium. We therefore propose a model in which the high levels of *Ttc21b* gene expression “load” the cells destined to make up the forebrain with TTC21B protein. Thus, we propose the TTC21B protein is present long after gene expression is diminished and is an especially perdurant protein compared to average protein half-lives (Belle et al. 2006; Price et al. 2010; Cambridge et al. 2011).

### Neuroprogenitor proliferation and differentiation is altered in Ttc21b-aln mutants

We measured rates of progenitor proliferation with immunohistochemistry for the mitotic marker pHH3 at multiple stages of forebrain development in *Ttc21b*^*aln*^ mutants. We noted an increased proportion of cells positive for pHH3 at E10.5 (Fig. 3 A-C). At E12.5, we do not see a change in the mitotic index (Fig. 3D-F) and by E14.5, we note a decrease in the number of proliferating cells (Fig. 3G-I). These data are consistent with the histological analysis we present where the proliferative ventricular zone is increased in mutants relative to wild-type at early stages in neurogenesis. The TBR2-positive intermediate progenitors are also a critical population for generating the proper brain size and are the result of asymmetric divisions of apical progenitors. We note a small decrease in the relative number of TBR2-positive cells at E12.5 (Fig. 3 J-L) but a marked increase at E14.5 (Fig. 3M-O). We also analyzed patterns of cell death but see no appreciable signal in control or *Ttc21b*^*aln*^ sections (Fig. S1. J-L).

**Figure 3.**
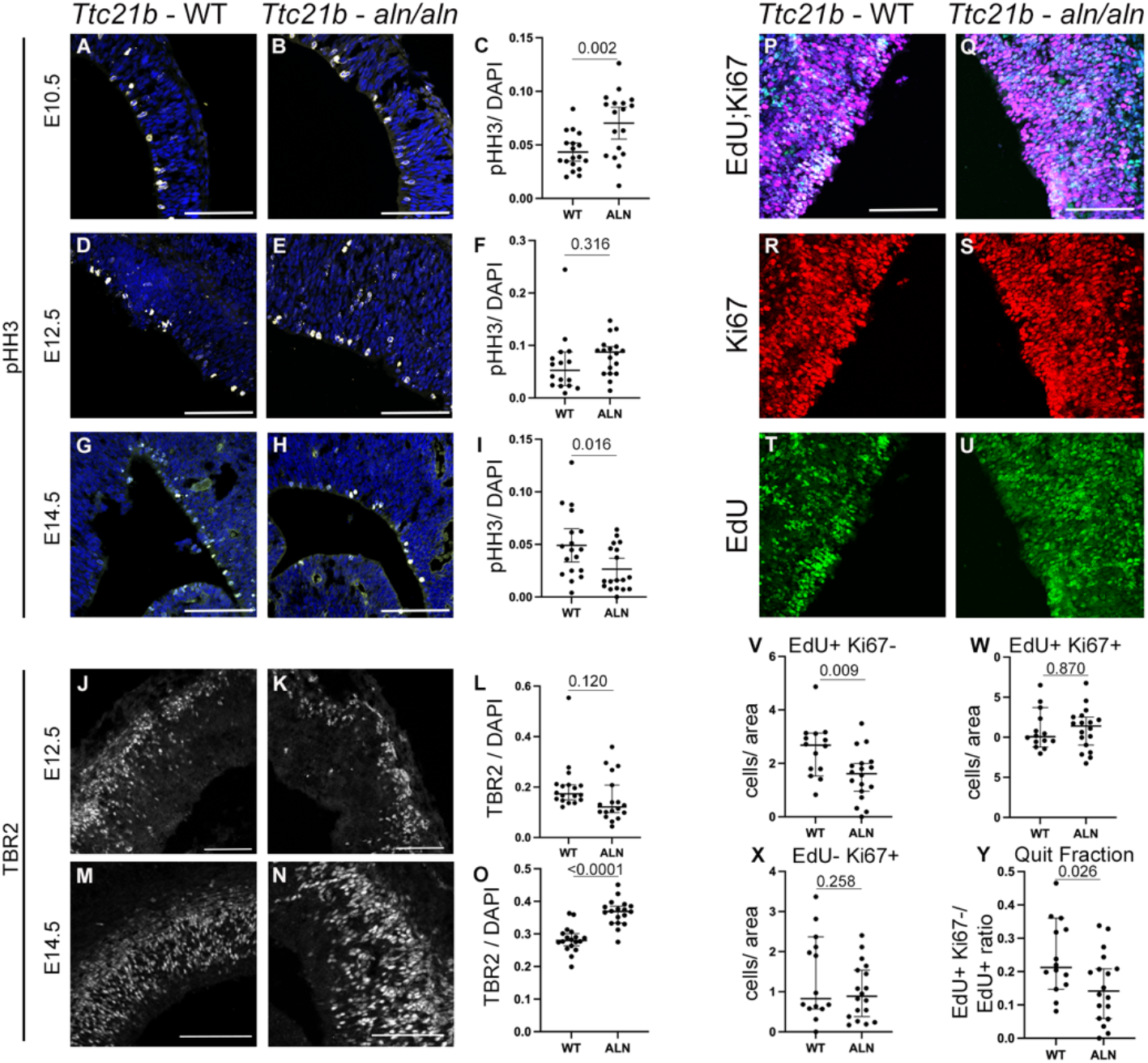
Neurogenesis in *Ttc21b*^*alien*^ cortical development. Immunohistochemistry for neuronal proliferation and differentiation markers was performed for pHH3 (A-I), TBR2 (J-O) and Edu/Ki67 (P-Y). Proliferation is marked by pHH3 at E10.5 in wild-type and *Ttc21b*^*aln/aln*^ mutants at E10.5 (A-C), E12.5 (D-F) and E14.5 (G-I). Quantifications of mitotic indices are shown for each age in C,F, I with t-test p values. TBR2-positive intermediate progenitors are shown at E12.5 (J-L) and E14.5 (M-O) and quantified in L,O. EdU and Ki67 immunohistochemistry are shown (P,Q, white cells are double labeled) with Ki67 (R,S) and EdU (T,U) shown individually. Quantifications are shown in V-Y as indicated in each. Scale bars = 100 µm.

**Figure 4.**
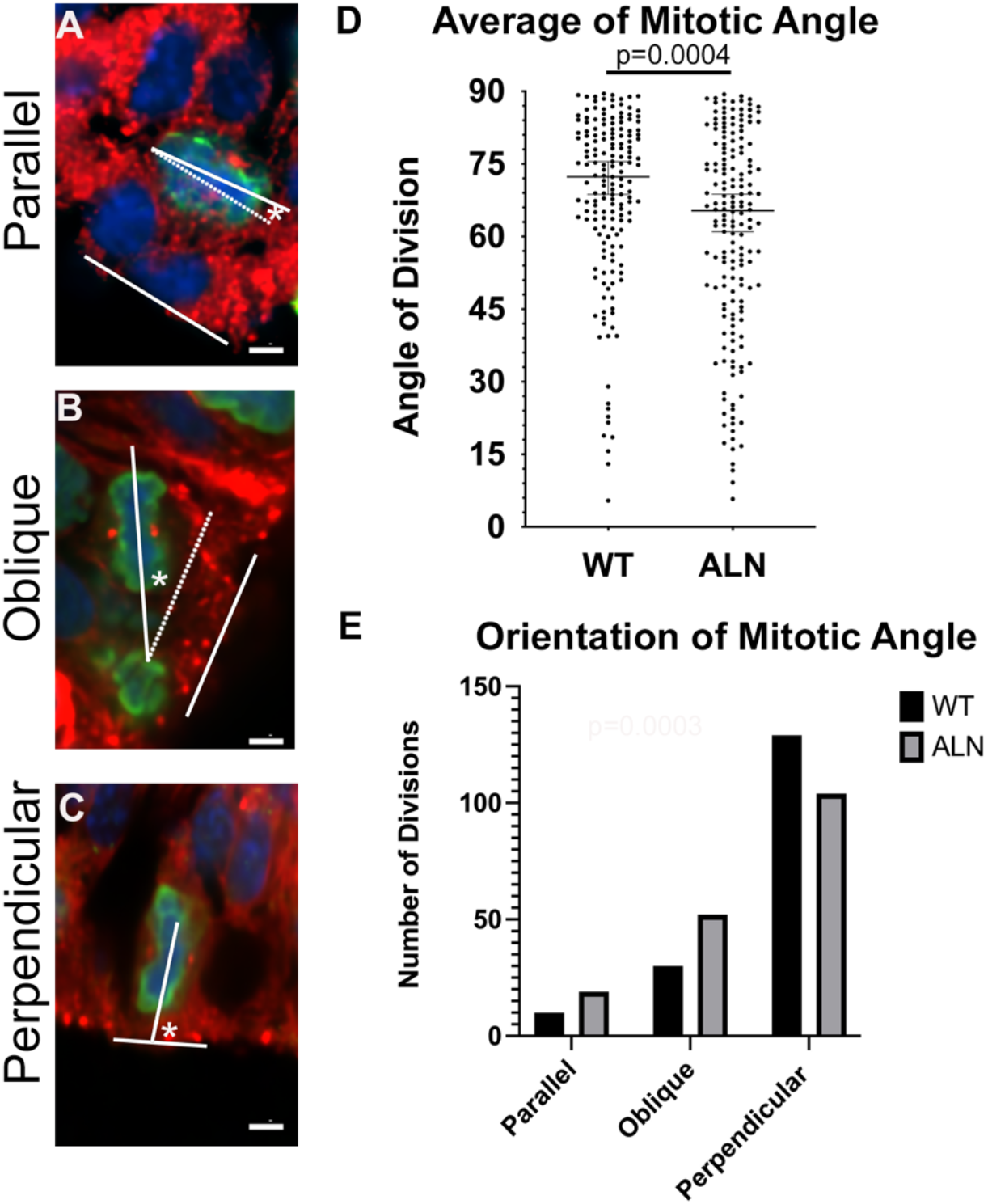
Mitotic angle of neural progenitors in *Ttc21b*^*alien*^ brain. The angle of the mitotic plane in dividing neural progenitors was determined with the use of immunohistochemistry for gamma tubulin and pHH3. Dividing cells were identified, a line was drawn identifying the plane of mitotic between the two centrosome, and a second line was drawn to identify the plane of the ventricular surface. The angle between these two lines is said to be the “mitotic angle.” Angles from 0°-30 ° are said to be parallel (example in A), 30 ° -60 ° are oblique (B) and 60 ° -90 ° are perpendicular (C). The average mitotic angle for individual mitoses wild-type and *Ttc21b*^*aln/aln*^ mutants are shown (D with t-test p value). The grouping of angles between parallel, oblique and perpendicular are shown in E. Scale bars = 10 µm.

In order to further understand patterns of proliferation in cells lacking *Ttc21b*, we labeled dividing cells with a maternal treatment of the thymidine analog EdU at E12.5 and examined embryos with immunohistochemistry at E13.5 for both EdU and the mitotic marker, Ki67 (Fig. 3P-U). This regimen allows analysis of cell behaviors over a 24 hour period of embryonic development. Cells that are positive for both EdU and Ki67 were actively dividing at both E12.5 (took up the EdU during DNA synthesis) and E13.5 (immuno-positive for Ki67). Those that are positive for EdU, but not Ki67 have not stayed in a proliferative mode, and this pool is commonly called the “quit fraction.” Quantification of these parameters shows that the number of EdU-positive dividing cells at E12.5 was somewhat reduced (Fig. 3V), and similar between controls and mutants at E13.5 (Fig. 3X). The number of double positive cells relative to area of the tissue was not changed (Fig. 3W). Interestingly, the EdU-positive, Ki67-negative cells as a proportion of EdU-positive cells (the quit fraction) was indeed reduced in mutants (Fig. 3Y). Thus, we conclude that neuroprogenitor dynamics are significantly altered in *Ttc21b*^*aln*^ mutant and an overactive proliferative pool as marked by pHH3 at E12.5 may not properly maintain the stem cell niche as fewer cells re-enter the cell cycle. This is consistent with the TBR2 experiments we report as a fraction of cells that do not re-enter cell cycle will differentiate into TBR2-positive intermediate progenitors.

### The mitotic angle of ventricular progenitors is slightly altered in Ttc21b^aln^ mutants

Given the behaviors we just described, we hypothesized the angle of the neuroprogenitor cell mitotic spindles relative to the plane of the ventricular zone may be different in *Ttc21b*^*aln*^ mutants. This mitotic angle has been previously correlated with the fate of the cells after mitosis (Taverna et al. 2014;

Matsuzaki and Shitamukai 2015). Cells with a plane of mitosis perpendicular to the plane of the ventricular zone are said to undergo a symmetric division with fairly equal division of cellular contents and are more likely to go on to generate two “mother” stem cells. In contrast, division planes more parallel to the VZ are thought to asymmetrically divide cellular contents and the cell closer to the VZ is likely to remain a stem cell while the “daughter” cell will begin to differentiate into a neuron, or a TBR2-positive basal progenitor. We used immunostaining for pHH3 (DNA) and gamma-tubulin to mark centrosomes for over 150 mitoses in both controls and mutants from multiple animals at E12.5. We measured the angle of mitosis (Fig. 5A-C) and found the *Ttc21b*^*aln*^ mutants had a smaller average angle of mitoses suggesting these were, on average, more asymmetric (Fig. 5D). We further subdivided these into three categories of parallel mitoses (angles 0-30°), oblique (30 ° -60 °) and perpendicular (60 ° -90 °). The mutants had slightly more parallel and oblique mitoses as compared to wild-type (Fig. 5E). All of these findings are consistent with the previous data and ultimate histological phenotypes described above and suggest that loss of *Ttc21b* leads to changes in centrosome dynamics in the proliferative neural epithelium.

**Figure 5.**
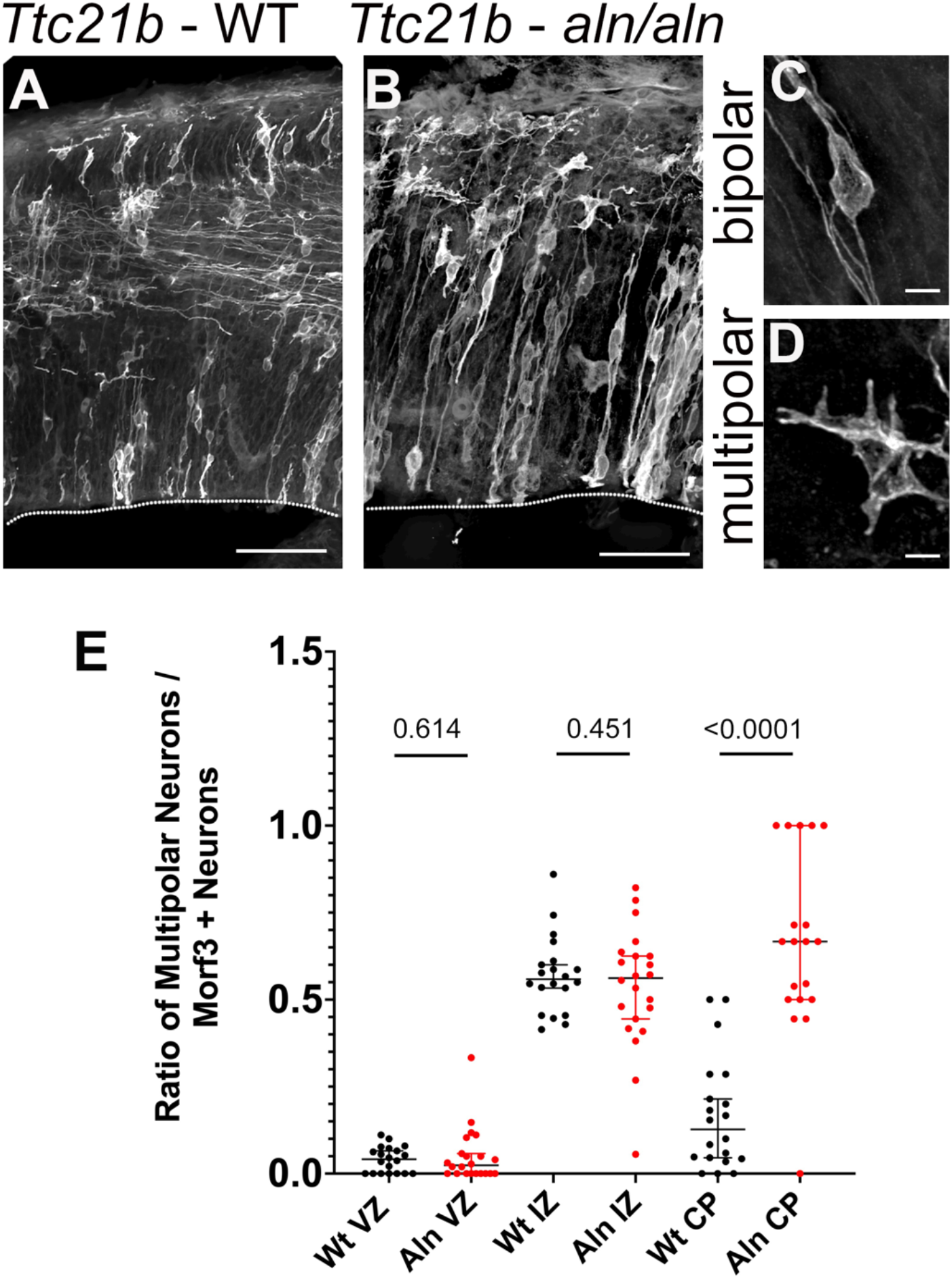
Cellular morphology in in *Ttc21b*^*alien*^ neurons. The MORF3 reporter allele was used with the *Emx1-Cre* line to stochastically label cells in the forebrain. Sections from WT (A) and *Ttc21b*^*aln/aln*^ mutant brains (B) show marked differences in cellular extensions made in the cortical plate (an example is highlighted by the arrow in A). Examples of bipolar (C) and multipolar (D) cells are shown and the ratio of these in each region of the brain are shown E for WT and *Ttc21b*^*aln/aln*^ mutants. The cortical plate quantification shows that *Ttc21b*^*aln/aln*^ mutant cells are much more likely to retain a multipolar morphology (E). Scale bars in A,B= 50 µm, C,D = 5 µm.

In order to further explore the latter stages of cortical development in the *Ttc21b*^*aln*^ mutants with the hypothesis that intracellular cytoskeletal dynamics are perturbed upon loss of *Ttc21b*, we incorporated the MORF3 reporter allele which will stochastically label individual clones within a Cre-positive lineage (Veldman et al. 2020). We used the *Emx1-Cre* to label some neurons within the developing forebrain in the control (*Emx1cre*/wt; *MORF3*/wt;*Ttc21b*^*wt/wt*^) and mutant (*Emx1cre*/wt; *MORF3*/wt;*Ttc21b*^*aln/aln*^). We first quantified the proportion of cells at E14.5 with a multipolar morphology (as compared to bipolar) in the VZ and intermediate zone (Fig. 5A-D). Radial migration includes the changes in cellular morphology from a multipolar arrangement to a bipolar arrangement as the migrating cell interacts with the radial glial cell scaffold used for the radial migration. Given the profound morphological deficits we see in the *Ttc21b*^*aln*^ mutants, we hypothesized these cellular reorganizations would be affected. Surprisingly, we did not see a significant difference in cellular morphologies at this multipolar to bipolar transition (Fig. 5 A,B,E). We did, however, see a striking change in cellular morphology as cells reach their final destination in the cortical plate as cells seemed to inappropriately stalled in a multipolar morphology (Fig. 5A,B).

## DISCUSSION

Here we report on a cellular basis for the striking microcephaly seen in the *Ttc21b*^alien^ homozygous mouse mutants. We show that early neural progenitor proliferation kinetics are disrupted upon loss of *Ttc21b* and see an accompanying shift in the angle of the mitotic spindle relative to the ventricular zone. We also note a change in the proportion of cells re-entering the cell cycle in mutant embryos. We show the surprising result that *Ttc21b* expression is not very prominent in the neural epithelium at the stages when these molecular phenotypes are observed. Rather, expression is much higher at earlier stages suggesting that TTC21B protein is especially perdurant and the relevant genetic ablations may be at significantly earlier stages. This would potentially explain why Cre transgenic lines commonly used in the field and expected to affect neural progenitor proliferation (i.e, *Foxg1-Cre* and *Emx1-Cre*; (Hebert and McConnell 2000; Gorski et al. 2002) did not recapitulate the null phenotype described here and previously reported (Stottmann et al. 2009).

Testing the model that the relevant *Ttc21b* expression domain is at earlier stages of tissues fated to become the telencephalon has been unexpectedly challenging. The *Hesx1-Cre* which has been shown to stimulate loxP recombination in the anterior neuroectoderm (Andoniadou et al. 2007) and would be a good candidate *Ttc21b* domain did not lead to robust recombination activity in our hands. We attempted an embryo-wide ablation of *Ttc21b* with an inducible Cre (*Gt(ROSA)26Sor*^*tm1(cre/ERT2)Tyj*^*/J*; (Ventura et al. 2007), but tamoxifen treatment at early stages (E4.5, E5.5) with a dose high enough to stimulate broad recombination lead to significant embryonic death precluding an efficient measure of brain size at late organogenesis stages. Direct measurement of TTC21B protein is hampered by current unavailability of robust antibodies. We constructed an epitope tagged allele of *Ttc21b* but this did not lead to effective translation of a tagged form TTC21B for reasons we have not been able to determine. Further testing of this model in future studies will necessitate the use of additional Cre transgenics and/or a new allele of *Ttc21b* with a degron tag to allow controlled degradation of endogenous protein.

Cell cycle defects in a cilia mutant such as those we highlight here are consistent as assembly and disassembly of the cilium is necessary to allow the centrosomes to be used in the mitotic apparatus. Several excellent reviews comprehensively discuss the literature in this area and we would particularly call attention to (Hasenpusch-Theil and Theil 2021; Liu et al. 2021; Zaidi et al. 2022). We hypothesized the dysmorphic cortex in the *Ttc21b*^*alien*^ homozygous mutants and previous evidence for a role for cilia in neuronal migration (Higginbotham et al.2012) would be the result of altered radial migration and an inability for the cell to properly employ cytoskeletal elements. We used the MORF3 allele to highlight cellular shape and similarly do not see evidence for a defect in the transition from a multipolar to bipolar morphology as neurons utilize the radial glial scaffolds for radial migration. However, we do see a marked decreased in cellular extensions in the cortical plate once cells do reach their final destination. Given that only some processes mediated by the cytoskeleton seem to be disrupted in *Ttc21b*^*alien*^ mutants, this may be a fruitful avenue for further investigation.

## ACKNOWLEDGEMENTS

This work was supported by the NIH (R35GM131875 to R.W.S.).

---

**Figure S1.**
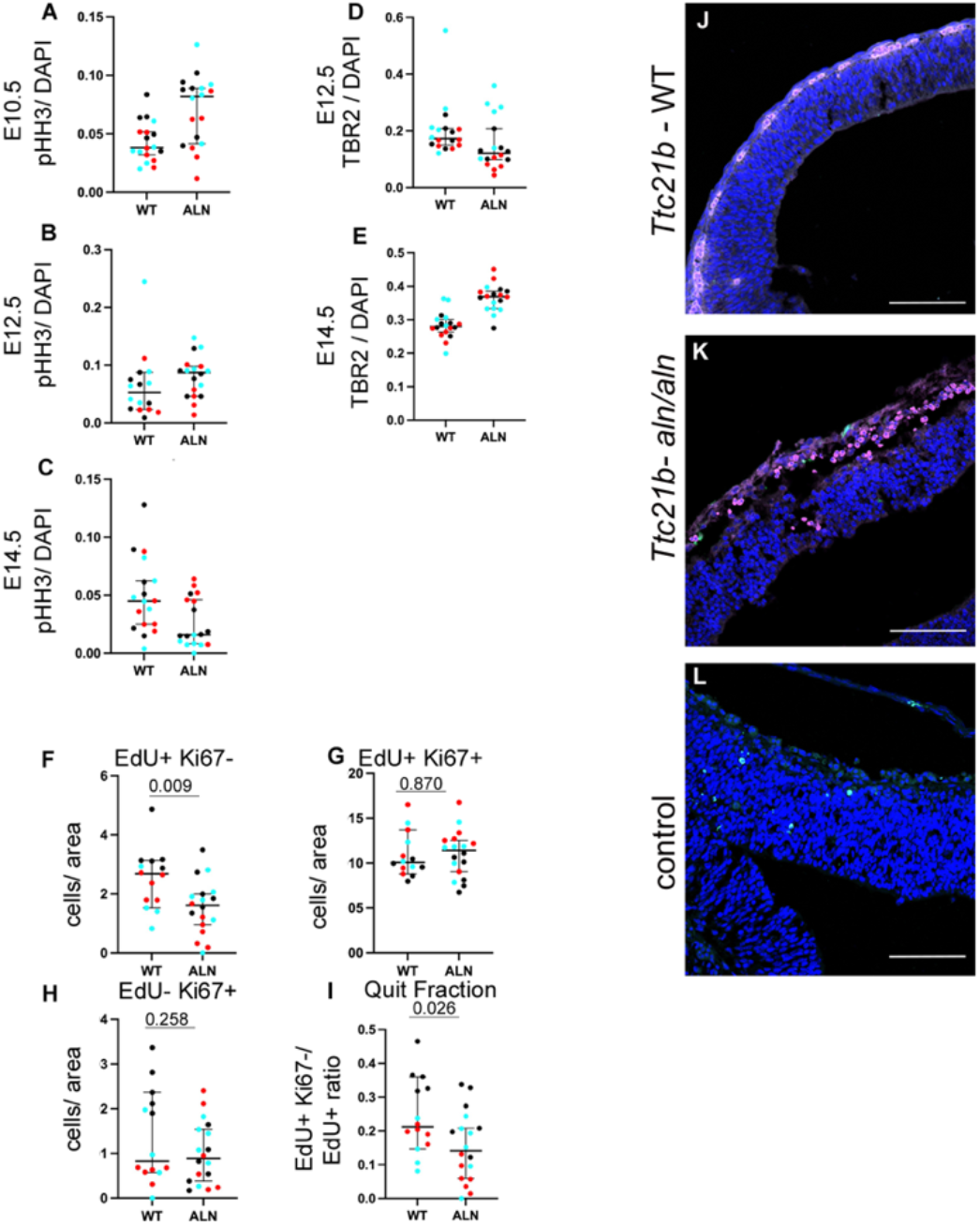
Neurogenesis in *Ttc21b*^*alien*^ cortical development. (A-I) Quantifications shown in Figure 3 C, F, I, L, O, V-Y are shown with different colors indicating which values from specific sections come from which individual embryos. The order of graphs shown in Figure 3 is repeated here. Cleaved-caspase 3 immunohistochemistry was used to assess cellular apoptosis in wild-type (J) and *Ttc21b*^*aln/aln*^ mutants (K) with no appreciable signal seen in each. A positive control from a different genotype (*Tubb2b*^*brdp*^, (Stottmann et al. 2013)) was used to validate the CC3 antibody (green cells in L are consistent with previous data showing apoptotic cells in this mutant (t-test p values shown). Scale bars = 100 µm.

## REFERENCES

Alberts B. 1994. Molecular biology of the cell. Garland Pub., New York.

Andoniadou CL, Signore M, Sajedi E, Gaston-Massuet C, Kelberman D, Burns AJ, Itasaki N, Dattani M, Martinez-Barbera JP. 2007. Lack of the murine homeobox gene Hesx1 leads to a posterior transformation of the anterior forebrain. Development 134: 1499–1508.

Baker K, Beales PL. 2009. Making sense of cilia in disease: the human ciliopathies. Am J Med Genet C Semin Med Genet 151C: 281–295.

Behringer R. 2014. Manipulating the mouse embryo : a laboratory manual. Cold Spring Harbor Laboratory Press, Cold Spring Harbor, New York.

Belle A, Tanay A, Bitincka L, Shamir R, O’Shea EK. 2006. Quantification of protein half-lives in the budding yeast proteome. Proc Natl Acad Sci U S A 103: 13004–13009.

Cambridge SB, Gnad F, Nguyen C, Bermejo JL, Kruger M, Mann M. 2011. Systems-wide proteomic analysis in mammalian cells reveals conserved, functional protein turnover. J Proteome Res 10: 5275–5284.

Goetz SC, Anderson KV. 2010. The primary cilium: a signalling centre during vertebrate development. Nat Rev Genet 11: 331–344.

Gorski JA, Talley T, Qiu M, Puelles L, Rubenstein JL, Jones KR. 2002. Cortical excitatory neurons and glia, but not GABAergic neurons, are produced in the Emx1-expressing lineage. J Neurosci 22: 6309–6314.

Hasenpusch-Theil K, Theil T. 2021. The Multifaceted Roles of Primary Cilia in the Development of the Cerebral Cortex. Front Cell Dev Biol 9: 630161.

Hebert JM, McConnell SK. 2000. Targeting of cre to the Foxg1 (BF-1) locus mediates loxP recombination in the telencephalon and other developing head structures. Dev Biol 222: 296–306.

Higginbotham H, Eom TY, Mariani LE, Bachleda A, Hirt J, Gukassyan V, Cusack CL, Lai C, Caspary T, Anton ES. 2012. Arl13b in primary cilia regulates the migration and placement of interneurons in the developing cerebral cortex. Dev Cell 23: 925–938.

Liu S, Trupiano MX, Simon J, Guo J, Anton ES. 2021. The essential role of primary cilia in cerebral cortical development and disorders. Curr Top Dev Biol 142: 99–146.

Matsuzaki F, Shitamukai A. 2015. Cell Division Modes and Cleavage Planes of Neural Progenitors during Mammalian Cortical Development. Cold Spring Harb Perspect Biol 7: a015719.

Price JC, Guan S, Burlingame A, Prusiner SB, Ghaemmaghami S. 2010. Analysis of proteome dynamics in the mouse brain. Proc Natl Acad Sci U S A 107: 14508–14513.

Reiter JF, Leroux MR. 2017. Genes and molecular pathways underpinning ciliopathies. Nat Rev Mol Cell Biol 18: 533–547.

Snedeker J, Gibbons WJ, Jr., Paulding DF, Abdelhamed Z, Prows DR, Stottmann RW. 2019. Gpr63 is a modifier of microcephaly in Ttc21b mouse mutants. PLoS Genet 15: e1008467.

Snedeker J, Schock EN, Struve JN, Chang CF, Cionni M, Tran PV, Brugmann SA, Stottmann RW. 2017. Unique spatiotemporal requirements for intraflagellar transport genes during forebrain development. PLoS One 12: e0173258.

Stottmann RW, Donlin M, Hafner A, Bernard A, Sinclair DA, Beier DR. 2013. A mutation in Tubb2b, a human polymicrogyria gene, leads to lethality and abnormal cortical development in the mouse. Hum Mol Genet 22: 4053–4063.

Stottmann RW, Tran PV, Turbe-Doan A, Beier DR. 2009. Ttc21b is required to restrict sonic hedgehog activity in the developing mouse forebrain. Dev Biol 335: 166–178.

Taverna E, Gotz M, Huttner WB. 2014. The cell biology of neurogenesis: toward an understanding of the development and evolution of the neocortex. Annu Rev Cell Dev Biol 30: 465–502.

Tran PV, Haycraft CJ, Besschetnova TY, Turbe-Doan A, Stottmann RW, Herron BJ, Chesebro AL, Qiu H, Scherz PJ, Shah JV et al. 2008. THM1 negatively modulates mouse sonic hedgehog signal transduction and afects retrograde intraflagellar transport in cilia. Nat Genet 40: 403–410.

Veldman MB, Park CS, Eyermann CM, Zhang JY, Zuniga-Sanchez E, Hirano AA, Daigle TL, Foster NN, Zhu M, Langfelder P et al. 2020. Brainwide Genetic Sparse Cell Labeling to Illuminate the Morphology of Neurons and Glia with Cre-Dependent MORF Mice. Neuron 108: 111–127 e116.

Ventura A, Kirsch DG, McLaughlin ME, Tuveson DA, Grimm J, Lintault L, Newman J, Reczek EE, Weissleder R, Jacks T. 2007. Restoration of p53 function leads to tumour regression in vivo. Nature 445: 661–665.

Wheway G, Nazlamova L, Hancock JT. 2018. Signaling through the Primary Cilium. Front Cell Dev Biol 6: 8.

Zaidi D, Chinnappa K, Francis F. 2022. Primary Cilia Influence Progenitor Function during Cortical Development. Cells 11.

